# Comparative Genomics and Integrated Network Approach Unveiled Undirected Phylogeny Patterns, Co-mutational Hotspots, Functional Crosstalk and Regulatory Interactions in SARS-CoV-2

**DOI:** 10.1101/2020.06.20.162560

**Authors:** Vipin Gupta, Shaiza Haider, Mansi Verma, Nirjara Singhvi, Kalaiarasan Ponnusamy, Md. Zubbair Malik, Helianthous Verma, Roshan Kumar, Utkarsh Sood, Princy Hira, Shiva Satija, Yogendra Singh, Rup Lal

**Author notes:** Contributed Equally (Vipin Gupta, Shaiza Haider, Mansi Verma). Author order was determined by drawing straws.

## Abstract

SARS-CoV-2 pandemic resulted in 92 million cases in a span of one year. The study focuses on understanding population specific variations attributing its high rate of infections in specific geographical regions particularly in USA. Rigorous phylogenomic network analysis of complete SARS-CoV-2 genomes (245) inferred five central clades named a (ancestral), b, c, d and e (subtype e1 & e2). The clade d & e2 were found exclusively comprising of USA. Clades were distinguished by 10 co-mutational combinations in Nsp3, ORF8, Nsp13, S, Nsp12, Nsp2 and Nsp6. Our analysis revealed that only 67.46% of SNP mutations were at amino acid level. T1103P mutation in Nsp3 was predicted to increase protein stability in 238 strains except 6 strains which were marked as ancestral type; whereas co-mutation (P409L & Y446C) in Nsp13 were found in 64 genomes from USA highlighting its 100% co-occurrence. Docking highlighted mutation (D614G) caused reduction in binding of Spike proteins with ACE2, but it also showed better interaction with TMPRSS2 receptor contributing to high transmissibility among USA strains. We also found host proteins, MYO5A, MYO5B, MYO5C had maximum interaction with viral proteins (N, S, M). Thus, blocking the internalization pathway by inhibiting MYO5 proteins which could be an effective target for COVID-19 treatment. The functional annotations of the HPI network were found to be closely associated with hypoxia and thrombotic conditions confirming the vulnerability and severity of infection. We also screened CpG islands in Nsp1 & N conferring ability of SARS-CoV-2 to enter and trigger ZAP activity inside host cell.

**Importance:** In the current study we presented a global view of mutational pattern observed in SARS-CoV-2 virus transmission. This provided a who-infect-whom geographical model since the early pandemic. This is hitherto the most comprehensive comparative genomics analysis of full-length genomes for co-mutations at different geographical regions specially in USA strains. Compositional structural biology results suggested that mutations have balance of contrary forces effect on pathogenicity suggesting only few mutations to effective at translation level but not all. Novel HPI analysis and CpG predictions elucidates the proof of concept of hypoxia and thrombotic conditions in several patients. Thus, the current study focuses the understanding of population specific variations attributing high rate of SARS-CoV-2 infections in specific geographical regions which may eventually be vital for the most severely affected countries and regions for sharp development of custom-made vindication strategies.

## Introduction

SARS-CoV-2 is a single stranded RNA virus with a genome size ranging from 29.8 kb to 29.9 kb (1). Most countries are facing the second waves and are on the verge of next wave. So far more than 18 million deaths and 800 million active cases have been reported worldwide (https://www.worldometers.info/coronavirus/). The genomic repertoire of SARS-CoV-2 comprises of 10 open reading frames (ORFs) encoding 27 proteins (2). ORF1ab encodes for 16 Non-structural proteins (Nsp) whereas structural proteins include spike (S), envelope (E), membrane (M), and nucleocapsid (N) proteins (3, 4). In addition, the genome of SARS-CoV-2 comprises of ORF3a, ORF6, ORF7a, ORF7b, ORF8 and ORF9 genes encoding six accessory proteins, flanked by 5’ and 3’ UTRs (1). In our previous study (5), a higher mutational rate in the genomes from different geographical locations around the world by accumulation of Single Nucleotide Polymorphisms (SNPs) was reported. Even during these early stages of the global pandemic, genomic surveillance has been used to differentiate circulating strains into distinct, geographically based lineages (6). However, the ongoing analysis of this global dataset suggests no consolidated significant links between SARS-CoV-2 genome sequence variability, virus transmissibility and disease severity.

Although there are several studies that have appeared ever since the emergence of SARS-Cov-2 (7, 8) and it has been reflected that the mutations at both genomic and protein level are in “Hormonical Orchestra” (9) that drives the evolutionary changes, demanding a detailed study of SARS-CoV-2 mutations to understand its successful invasion and infection. To unveil this, we rendered and screened 18775 genomes of SARS-CoV-2 and selected 245 genomic sequences deciphering the phylogenetic relationships, tracing them to SNPs at nucleotide and amino acid variation (AAV) levels and performing structural re-modelling. We specifically focused on the evolutionary relationships among the strains predicting Nsp3 as mutational hotspot for SARS-CoV-2. Study was extended to understand the mechanism of host immunity evasion by Host-Pathogen Interaction (HPI) and confirming their interactions with host proteins by docking studies. We identified sparsely distributed hubs which may interfere and control network stability as well as other communities/modules. This indicated the affinity to attract a large number of low-degree nodes toward each hub, which is a strong evidence of controlling the topological properties of the network by these few hubs (10). We also analyzed the transfer of genomic SNPs to amino acid levels and associations of CpG dinucleotides contributing towards the pathogenicity of SARS-CoV-2. Since the CpG islands have always been linked with epigenetic regulation and act as the hotspots for methylation in case of viruses (11–13). But for RNA viral genomes, CpG nucleotides are the targets for Zinc Antiviral protein (ZAP), a major factor of mammalian interferon-mediated immune response (14, 15). Here also, the conservancy found in possession of CpG dinucleotides towards the extremities of all the genomes considered in the present analysis indicate their importance in evading host immunity.

## Material and Methods

### Selection of genomes, annotations and phylogeny construction

Publicly available genomes of SARS-CoV-2 viruses were obtained from the NCBI database (https://www.ncbi.nlm.nih.gov/genbank/sars-cov-2-seqs/). Until March 31, 2020 only 447 (Data Set S1, Sheet1) SARS-CoV-2 genomes were available in the databases (Supplementary data). The data was screened for unwanted ambiguous bases using N-analysis program, based on which 245 (Data Set S1, Shee2) complete and clean genomes of SARS-CoV-2 were selected for further analysis (Supplementary data). A manually annotated reference database was generated using GenBank file of severe acute respiratory syndrome coronavirus 2 isolate SARS-CoV-2/SH01/human/2020/CHN (Accession number: MT121215.1) and open reading frames (ORFs) were predicted against the formatted database using prokka (−gcode 1) (16). Genomic sequences included in the analysis belongs to different countries namely, USA (168), China (53), Pakistan (2), Australia (1), Brazil (1), Finland (1), India (2), Israel (2), Japan (5), Vietnam (2), Nepal (1), Peru (1), South Korea (1), Spain (1), Sweden (1). Whole genomes nucleotide and protein sequences were aligned using mafft (17) at 1000 iterations. The alignments so obtained were processed for phylogeny construction using BioEdit software (18). The nucleotide-based phylogeny was annotated and visualized on iTOL server (19). While amino acid-based phylogeny was visualized and annotated using GrapeTree (20).

### Genotyping based on SNP/AAV

To detect nucleotide and amino acid variations (AAV) among 245 genomes of SARS-CoV-2, sequence alignment of nucleotide and amino acid, respectively were performed against the reference genome. The change of nucleotide and amino acid was calculated as point variations and were recorded. The interpolation and visualization were plotted using computer programs in Python. Co-mutation were predicted, and clustering was performed using MicroReact (21). For validation we selected 18775 (Data Set S1, Sheet3) complete genomes available NCBI virus database (https://www.ncbi.nlm.nih.gov/labs/virus/vssi/#/virus. Last accessed in September 2020. After removing the genomes containing sequencing errors and unidentified base pairs “N”, remaining 12299 genomes were used (Data Set S1, Sheet4).

### Data and Computer programs

The genomic analytics is performed using programs in Python and Biopython libraries (22). The computer programs and the updated SNPs profiles of SARS-CoV-2 isolates are available upon requests.

### Construction of the Host-Pathogen Interaction Network of SARS-CoV-2

The interactions between viral and host proteins are responsible for all aspects of the viral life cycle; from infection of the host cell, to replication of the viral genome, and assembly of new viral particles (23). To find the Host Pathogen Interaction (HPI), we subjected SARS-CoV-2 proteins sequence to Host-Pathogen interaction databases such as Viruses STRING v10.5 (24) and HPIDB3.0 (25) to predict their direct interaction with human as the principal host. In these databases, the virus–host interaction was imported from different PPI databases like MintAct (26), IntAct (26), HPIDB (25) and VirusMentha (27). It searches protein sequences using BLASTP to retrieve homologous host/pathogen sequences. For high-throughput analysis, it searches multiple protein sequences at a time using BLASTp and obtain results in tabular and sequence alignment formats (28). The HPI network was constructed and visualized using Cytoscape v3.7.2 (29). It is an open-source software platform for visualizing molecular interaction networks which involve various biological pathways and integrating these networks with annotations, gene expression profiles and other state data. In the constructed Network, proteins with highest degree, which interact with several other signaling proteins in the network indicate a key regulatory role as a hub. In our study, using Network Analyzer (30), plugin of Cytoscape v3.7.2, we identified the hub protein. Further, the human proteins interacting with individual viral proteins were subjected to functional annotation. Gene ontology (GO) analysis was performed using ClueGo (31), selecting the Kyoto Encyclopedia of Genes and Genomes (KEGG) (32), Gene Ontology—biological function database, and Reactome Pathways (33) databases. The ClueGo parameters were as follows: Go Term Fusion selected; pathways or terms of the associated genes, ranked based on the P value corrected with Bonferroni stepdown (P values of <0.05); GO tree interval, all levels; GO term minimum number of genes, 3; threshold, 4% of genes per pathway; kappa score, 0.42. Gene ontology terms are presented as nodes and clustered together based on the similarity of genes corresponding to each term or pathway.

### Computational structural analysis on wild-type and mutant SARS-CoV-2 proteins

SARS-CoV-2 proteins sequences were retrieved from the NCBI genome database and pairwise sequence alignment of wild-type and mutant proteins were carried out by the Clustal Omega tool (34). The wild-type and mutant homology model of S-protein, Nsp12 and Nsp13 were constructed using the SWISSMODEL (35), whereas the 3D structure of ORF8, ORF3A, Nsp2, Nsp3 and Nsp6 were predicted using Phyre2 server (36). The crucial host proteins (TMPRSS2, RPS6, ATP6V1G1 and MYO5C) 3D structures were generated using the SWISSMODEL and ACE2 structure retrieved from the PDB database (PDB ID: 6M17). These structures were energy minimized by the Chiron energy minimization server (37). The effect of the mutation was analyzed using HOPE (38) and I-mutant (39). The I-mutant method allows us to predict the stability of the protein due to mutation. The docking studies for wild and mutant SARS-CoV-2 proteins with host proteins was carried out using PatchDock Server (40). Structural visualizations and analysis were carried out using pyMOL2.3.5 (41).

### Analysis of CpG regions

SARS-CoV-2 genomes were analyzed for the presence of CpG regions that can be targeted for methylation induced gene silencing. To locate the CpG regions, meth primer 2.0 (http://www.urogene.org/methprimer2/) and the CpG Plot (http://www.ebi.ac.uk/Tools/emboss/cpgplot/) programs were used, although some variations were found in both the programs. Both the programs were run on default parameters of a sequence window longer than 100 bp; GC content of ≥50%, and an observed/expected CpG dinucleotide ratio ≥0.60. The presence of common CpG islands was confirmed by performing BLAST using the above reference strain.

## Results and Discussion

### Phylogenetic relationship between different SARS-CoV-2 strains

In our previous study, we reported a mosaic pattern of phylogenetic clustering of 95 genomes of SARS-COV-2 isolated from different geographical locations (5). Strains belonging to one country were found clustered with distant countries strains but not with the neighboring one. Taking clue from this study we constructed phylogenetic relatedness of 245 strains of SARS-COV-2 from USA, China, and several other countries including, Spain, Vietnam, Peru, Finland and Pakistan and unravel the significant association of evolutionary patterns among SARS-CoV-2 based on their geographical locations predicting their mosaic phylogenetic arrangements. It was found that most strains from USA were clustered together, but comparatively high divergences were found in strains isolated from China and Japan. Japanese strains were found to be scattered and formed clusters with strains from USA, Pakistan, Vietnam, Taiwan, and China. Even with a smaller number of genomes sequences from Japan, Vietnam and Peru revealed a highly scattered pattern and close associations with that of USA and Chinese strains were revealed. Strains reported from patients of Taiwan (MT192759), Australia (MT007544), South Korea (MT039890), Nepal (MT072688) and Vietnam (MT192773, MT192772) had travel histories from Wuhan, China (42). However, a strain from Pakistan (MT240479) which clustered with the Japanese strains was found to be isolated from patient having travel history from Iran. Indian strains (MT050439, MT012098) that were isolated from patients who travelled from Dubai, clustered with Chinese strains. Later, reports confirmed many cases of SARS-CoV-2 in Dubai from China (https://www.newsbytesapp.com/timeline/India/58169/271167/coronavirus-2-positive-cases-detected-in-delhi-telangana). Thus, a clear landscape of phylogenetic relationships could be obtained reflecting mosaic clustering patterns in accordance with the travel history of patients (Figure 1A). However, results were in contradiction with the genomic analysis of SARS-COV-2 by Forster et al., 2020 where they predicted the linear/directive evolution from ancestral node a to node b and c. Whereas we report here both divergent (from ancestral node a to b, c & e) and directive (**node c to d**) evolution among the SARS-CoV-2 strains (Figure1B).

**Figure 1.**
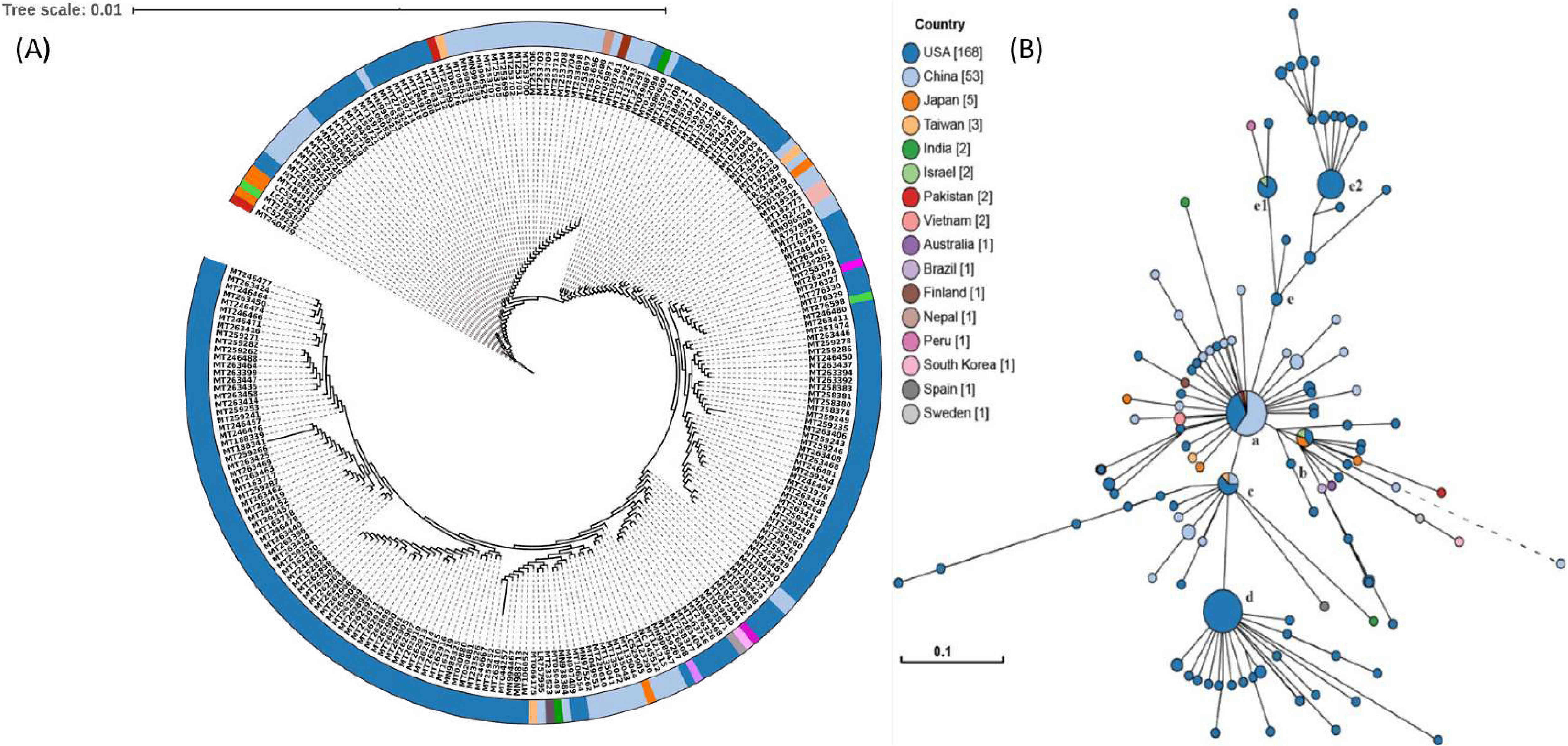
Phylogenetic network of 245 SARS-CoV-2 genomes. (A) Nucleotide based phylogenetic analysis of SARS-CoV-2 isolates using the Maximum Likelihood method based on the Tamura-Nei model, (B) Amino acid based phylogenomic analysis. Circle areas are proportional to the number of taxa. The map is diverged into 5 major clade (a-e) representing variation in the genomes at amino-acid level. The colored circle represents the country of origin of each isolate.

Since genome-based phylogeny did not highlighted the amino acid level changes, thus to ascertain the variations among the SARS-CoV-2 strains at protein level, we constructed whole proteome alignment-based phylogeny, clustered the 245 strains into five major **clades a-e (**Figure 1B). The first cluster, Clade-a had maximum nodes (46), including reference node, and strains from Nepal (MT072688), Pakistan (MT262993), Taiwan (MT192759) along with 15 strains from USA and 27 strains from China. It also had the mutated daughter nodes radiating outwards, belonging to China, Finland (MT020781), India (MT012098), Japan (LC534419, LC529905), Taiwan (MT066176), Vietnam (MT192772-3), Brazil (MT126808), Australia (MT007544), South Korea (MT039890) and Sweden (MT093571) along with seven USA strains (Figure 1B). This clade represented the ancestral node as it harbored the oldest known SARS-CoV-2 strain from China and laid the foundation of rest of the mutated daughter strains worldwide, marking the onset of the divergence in SARS CoV-2. Three significantly diverged network nodes originated from the ancestral clade-a and were marked as clade-b, c and e (Figure 1B). For **Clade-b**, central node included only four strains in which two were from USA (MT184912, MT276328) and one each from Israel (MT276597) and Japan (LC528233). Its major descended radiant belonged to Japan (LC528232, LC534418), Pakistan (MT240479), USA (MT184913, MT184910, MN997409) and China (MT049951, MT226610). It was observed that one of the Chinese strains in clade-b (MT226610) had the longest branch length making the strain very distinct (harboring 25 other mutations) by showing exceptionally high rate of evolution. In **Clade-c** lineage, small central node was comprised of Taiwan (MT066175), USA (MT246667, MT233526, MT020881, MT985325, MT020880) and Chinese (MN938384, LR757995) strains. Interestingly one strain each from Spain (MT233523) and India (MT050493) were also found radiating as daughter node from the central one. **Clade-d** lineage, which was originated from clade-c lineage, consisted only of USA strains both in central nodes and radiations. Importantly, 2 strains (MT263416, MT246471) were found most divergent with varied mutation suggesting the high rate of evolution among USA strains which might be linked with the high pathogenicity among them. **Clade-e** bifurcated into two sub-clads (e1 and e2) by significant set of mutations. Sub-clad-e1 include six strains from USA, one from Israel (MT276598) with radiating nodes from Peru (MT263074) and USA (MT276327); whereas sub-clad e2 had 32 strains belonging to USA. Effect of amino acid mutations were further checked on another subset of 12299 SARS-CoV-2 genomes (screened from 18775) for the validation. The random explosion of evolutionary clades were seen (Supplementary figure 1). There were other nodes progressing from e (e1-e2) to f (exclusive USA strains), g (g1), h, i, j (exclusive Australian strains) and k sub-clades. This divergence supported the random evolution of SARS-CoV-2 suggesting network expansion in multiple clades contradicting to the earlier directed evolution proposed by Forster et al., 2020. Also, the mutational counts (Data Set S3) observed by 12299 genomes were almost similar to those identified in 245 representative genomes (Supplementary figure 1). Thus, formation of five major evolutionary clades and subclades based on the amino acid phylogeny needs attention for identifying the assessment of divergence among SARS-CoV-2 strains.

### Genotyping and variation estimation

To understand the implication of mosaic pattern of transmissions and evolutionary lineage clustering (Clade a-e), we studied the SNPs genotyping from the 245 genome sequences as mutation counts along with their frequency at specific genomic locations. Mutational changes at protein/amino acid levels were also weighed by assessing AAV. Interpolations of the SNPs/AAVs data were made by assessing their frequency, genomic positions, and type of SNPs/AAVs (Figure 2B), highlighted a large mutational diversity among the virus isolates. We identified a total of 12 SNP types (A>G, A>C, A>T, C>A, C>G, C>T, G>A, G>C, G>T, T>A, T>C, T>G) accounting for mutations at 297 genomic locations (Figure 2A, 2B,). Overall pattern of SNPs suggested C>T transition as the most common mutation in the entire genomic sets (Figure 2A), however highest frequency was recorded for T>C transitions (Figure 2B). Based on the genomic arbitrators SNP frequencies, we analyzed 14 major locations inside the genomes of SARS-CoV-2 for potential mutation generating different allelic forms for genes (Table 1). The SNP of C>T was first observed at 67^th^ location in 5’ UTR region of leader sequence with a frequency of 45 followed by Nsp2 at two locations (885 & 2863) with the frequency of 29 and 44, respectively. Nsp3/PL-PRO and Nsp8 marked the highest frequency of 238 SNP counts of T>C at 5852 and 12299 locations. Another T>C SNP was observed in ORF8 with frequency of 88 at 27973 location. C>T SNP transformation was found in Nsp4 and Nsp12 with the frequency of 88 and 44 at location 8608 and 14234, respectively. Non-structural protein, Nsp13 was strangely found harboring two different SNP (C>T and A>G) at three different locations (17573, 17684, 17886, Figure 2 B) with a relatively high frequency of 68, 63 and 63 respectively. A>G SNP conversion in S (Spike) protein was found with a frequency of 43. A Low SNP count of G>T transitions were falling in the ORF3a and Nsp6 with frequency of 32 and 21, respectively (Table 1). Though, all SNP counts do not reflect the change at protein level and therefore must be estimated at the translation levels for their significant effect. Although 297 genomic locations harbored SNPs but their corresponding AAV were found only in 200 genomic locations accounting for 67.34% conversion efficiency. Out of 14 high frequency SNPs, only 9 mutations [Nsp2 (T85I), Nsp3 (S1103P), Nsp6 (L37F), Nsp12 (P324L), Nsp13 (P409L, Y446C), S (D614G), Orf3a (Q577H), Orf8 (L84S)] were found to reflect at protein level with the highest frequency of 238 in Nsp3 (Table 1).

**Table 1:**
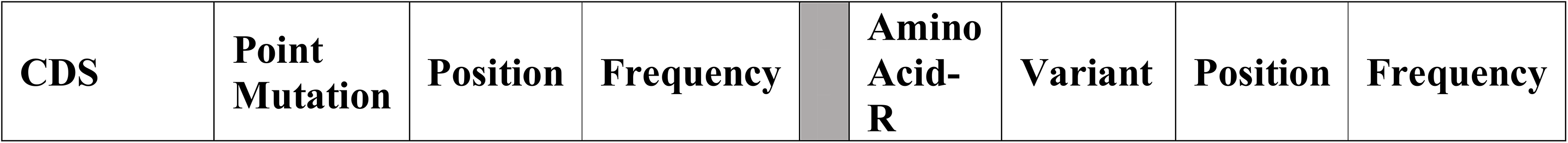
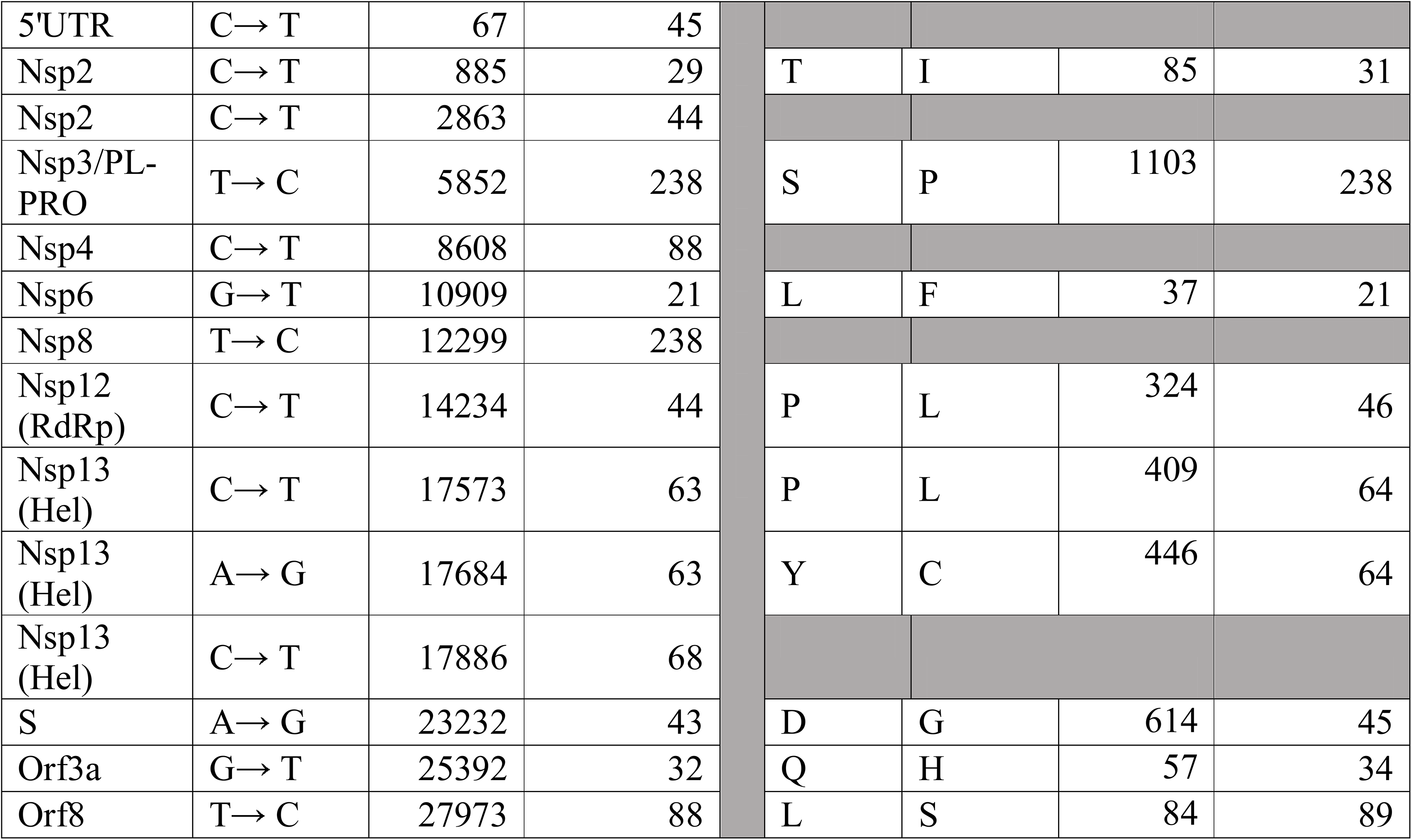
Common SNP and AAV mutations occurring in SARS CoV-2 genomes

**Figure 2.**
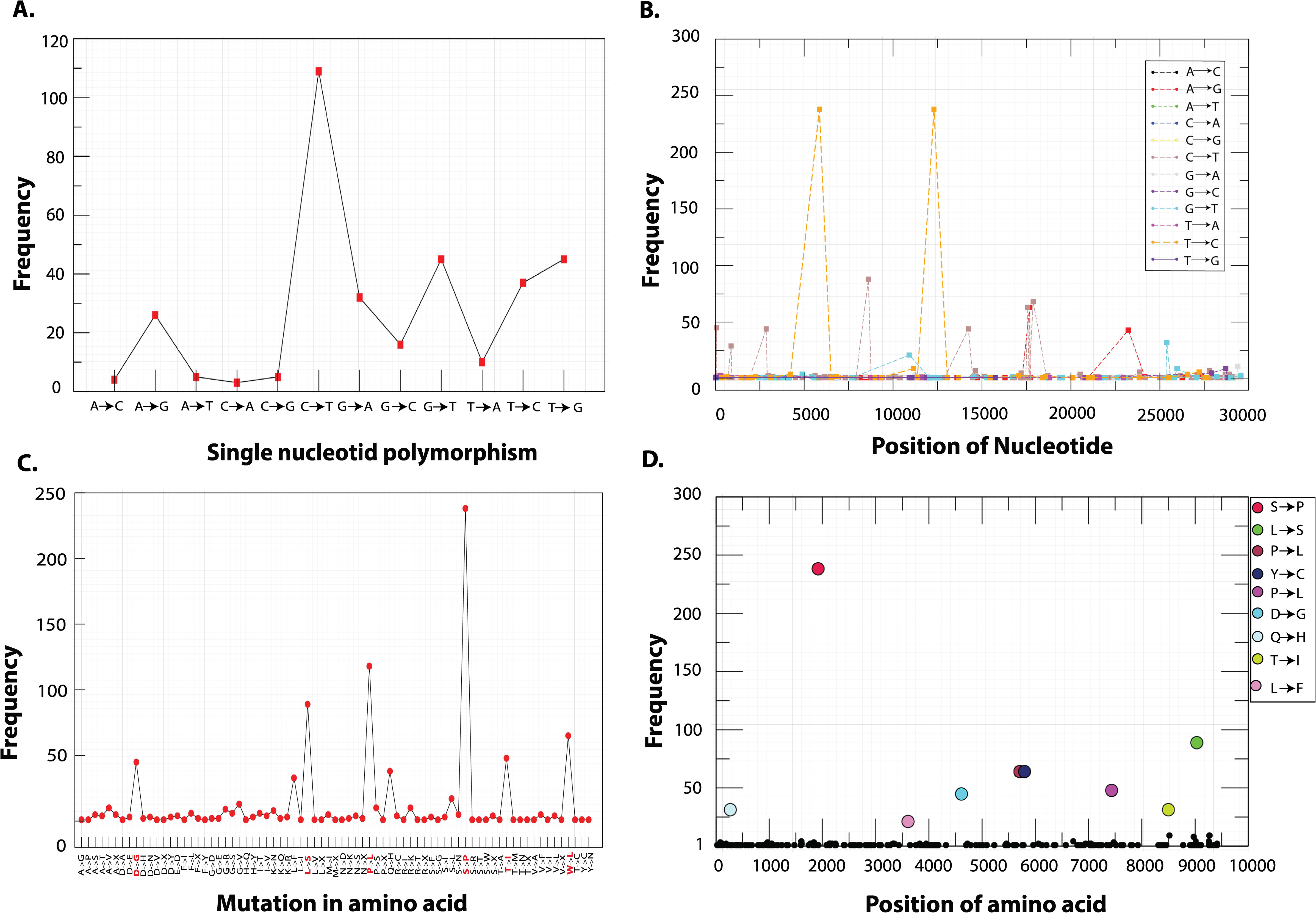
Distribution of SNP (A, B) and AAV (C, D) mutations of SARS-CoV-2 isolates from the globe. (A) Frequency based plot of 12 possible SNP mutations across 245 genomes, (B) Frequencies of the single SNP mutations with locations on the genome, (C) AAV based mutations across the genomes, (D) Top 9 AAV mutations holding highest frequencies among 245 genomes and their respective positions. The nucleotide and amino-acid positions are based on the reference genome of SARS-CoV-2.

These mutated proteins are known to play various regulatory roles and therefore, mutations at amino acid level can modulate their catalytic activity drastically. Specifically, Nsp3 is the largest and essential component of replication complex in the SARS-CoV-2 genome (43) and along with Nsp2 it forms a transcriptional complex in endosome of the infected host cell (44). Nsp6 is a multiple-spanning transmembrane protein located into the endoplasmic reticulum where they induce autophagosomes via an omegasome intermediate (45). Interestingly, the mutation of L37F caused stiffness in the secondary structure of Nsp6 and leads to low stability of the protein structure as observed in most recent strains isolated from Asia, America, Oceania and Europe (46). Nsp12 and Nsp13 are the key replicative enzymes, which require Nsp6, Nsp7 and Nsp10 as cofactors. Nsp12, RNA dependent RNA polymerase (RdRp) with the presence of the bulkier leucine side chain at location 324 is likely to create a greater stringency for base pairing to the templating nucleotide, thus modulating polymerase fidelity (47). Nsp13 contains a helicase domain, allowing efficient strand separation of extended regions of double-stranded RNA and DNA (48). Dual mutations in Nsp13 were reported with profound effect on its activity specifically in Pacific northwest of USA (49). P409L, mutation leads to increased affinity of helicase RNA interaction, whereas Y446C is a destabilizing mutation increasing the molecular flexibility and leading to decreased affinity of helicase binding with RNA (50). Therefore, both the mutations were antagonistic in nature. Thus, ORF1ab polyprotein of SARS-CoV-2 encompasses mutational spectra where signature mutations for Nsp2, Nsp3, Nsp6, Nsp12 and Nsp13 have been predicted.

Amino acid mutations in structural proteins S, ORF3a and ORF8 have also been observed with varied frequency of 45, 34 and 89 respectively. The mutation in Spike protein (D614G) has been reported to outcompete other preexisting subtypes, including the ancestral one. This mutation generates an additional serine protease (Elastase) cleavage site in Spike protein (51) which is discussed in more details in later sections. ORF3a mutation (Q57H), is located near TNF receptor associated factor-3 (TRAF-3) regions and has been reported as molecular difference marker in many genomes including Indian SARS-CoV-2 genomes (52) for their delineation. Mutation in ORF8 sequence (L84S) was found conserved (53) therefore to predict its effect it was critical to examine its biological function in SARS-CoV-2 interaction with human proteins.

Our results showed that the mutations (SNPs and AAV) in the virus were not uniformly distributed. Genotyping study annotated few mutations in the SARS-CoV-2 genomes at certain specific locations with high frequency predicting their high selective pressure. Thus, mutations can be predicted as location-specific but not type-specific by SNP count. Highly frequent AAV might be associated with the changes in transmissibility and virulence behavior of the SARS-CoV-2. Therefore, high-frequency AAV mutations in Spike protein, RdRp, helicase and ORF3a are important factors to consider while developing vaccines against the fast-evolving strains of SARS-CoV-2.

### Prevalence of Co-mutation in SARS-COV-2 evolution

Interestingly, we observed co-mutations in Nsp13 at locations 446 (Nsp13_1) and 409(Nsp13_2) that were prevalent in common 64 genomes, all belonging to USA. The AAV reported above (Table 1) were further analyzed and found occurring in 10 different permutations varying from single to multiple mutated protein combinations. Complete details of these co-mutations combinations are given in Table 2 Data Set S2. These co-mutations were mapped over the divergent phylogeny for indicating the evolutionary divergence among the 245 strains. The phylogram (Figure 1B) showed clear divergence of strains from the parent strain due to accumulation of mutations at different level of human-to-human transmission. We found co-mutations in Nsp3, ORF8, Nsp13, S, Nsp12, Nsp2 and Nsp6 were responsible for the above divergence.

**Table 2:**
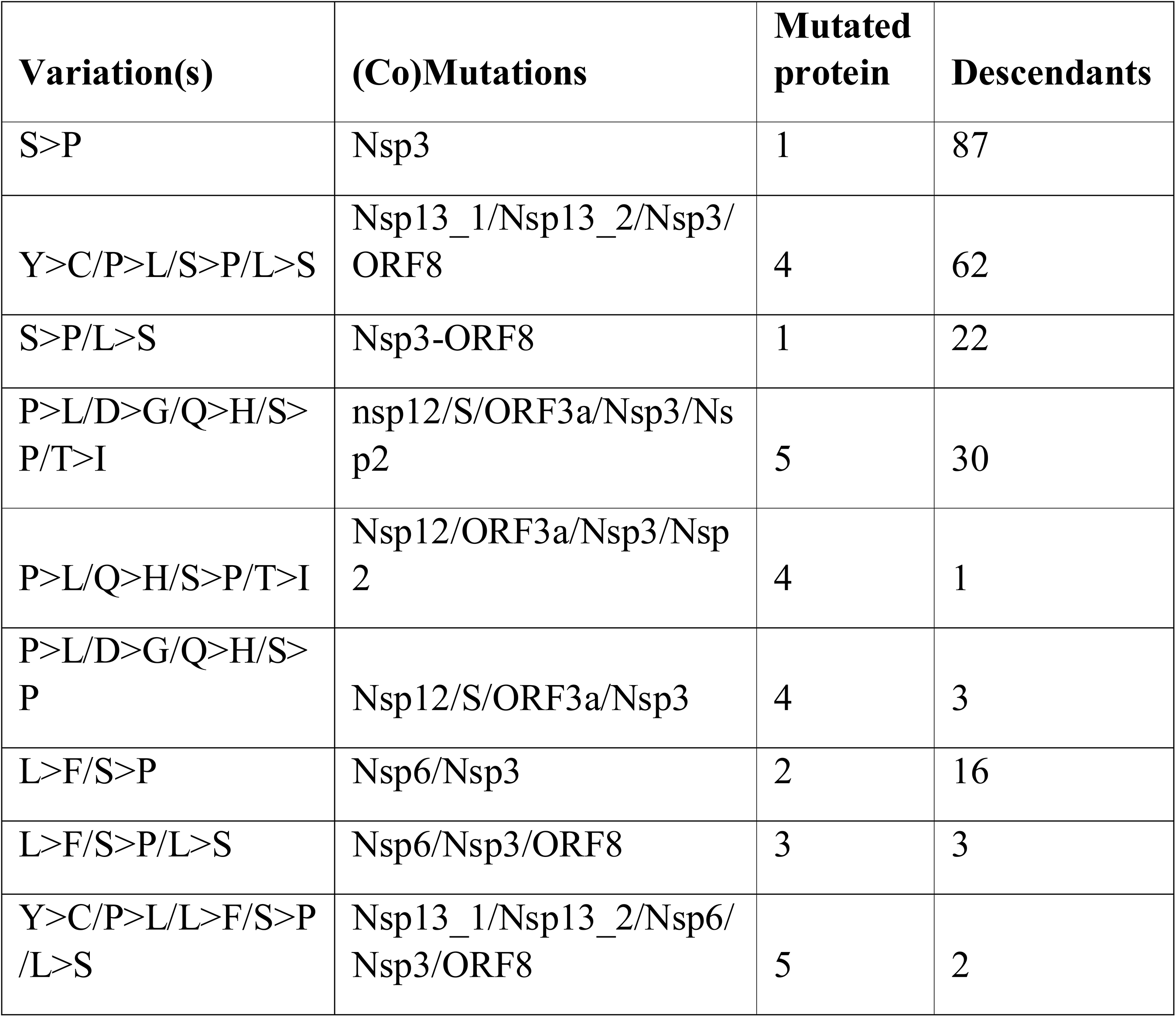
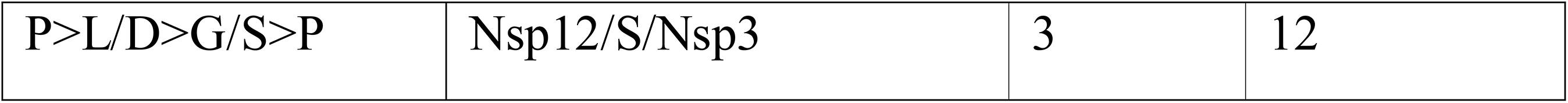
Co-mutations combinations and genomic location identified in different proteins of SARS-COV-2

These co-mutations were found linked with lineage **clades a to e,** highlighting their prevalence of delineation among them (Figure.1B). In **clade-a,** 40 genomes harbored mutations at only Nsp3 protein while six isolates belonging to USA (MT262993, MT044258, MT159716, MT259248, MT259267) and Pakistan (MT263424) showed no mutation confirming their lineage same as that of the reference/ancestral genome from China. Presence of Nsp3 mutation (S1103P) in 238 strains underlined the origin of mutation from reference strain highlighting the first mutational induced divergence in SARS-CoV-2 strains. Therefore, Nsp3 was marked as first mutational hotspot for accumulating amino acid mutations in SARS-CoV-2. Strains from Brazil (MT126808) and USA (MT276331) form the descendent from **clade-a** harboring Nsp3/Nsp6 as first mutational combination directing the common evolutionary lineages. The **clade-b** had an additional mutation of ORF8 along with Nsp3 and Nsp6 with three descendant strain from USA and China. We observed most distant Chinese strain (MT226610), clustered in **clade-b** and harbored additional 25 AAV making it the highly pathogenic strain in the network (Figure 1B). The **clade-c** descended from **clade-a** had a different set of co-mutation with Nsp3-ORF8 proteins while **clade-d** descended further from **clade-c** had two mutation in Nsp13 (P409L/Y446C) in addition to Nsp3/ORF8 proteins. Two strains from USA in the cluster radiating from **clade-d** harbored additional Nsp6 mutation stating them more divergent with scope of further possible evolution. The next subclade-e1 was found holding another new set of co-mutation of Nsp3/S/Nsp12. Whereas the highest number of co-mutations were found in subclade-e2 with combination of Nsp3/Nsp2/Nsp12/S/ORF3a prevalent in 30 genomes belonging to USA predicting them as active carrier of evolutionary force for SARS-CoV-2 divergence (Figure 3). In future, addition of more and more genome may indicate the evolutionary relationships among these co-mutations. Our result suggested that co-mutations are the major evolutionary force that drives the pathogenicity among the different geographical isolated strains which can be responsible for higher and lower order of virulence among them.

**Figure 3.**
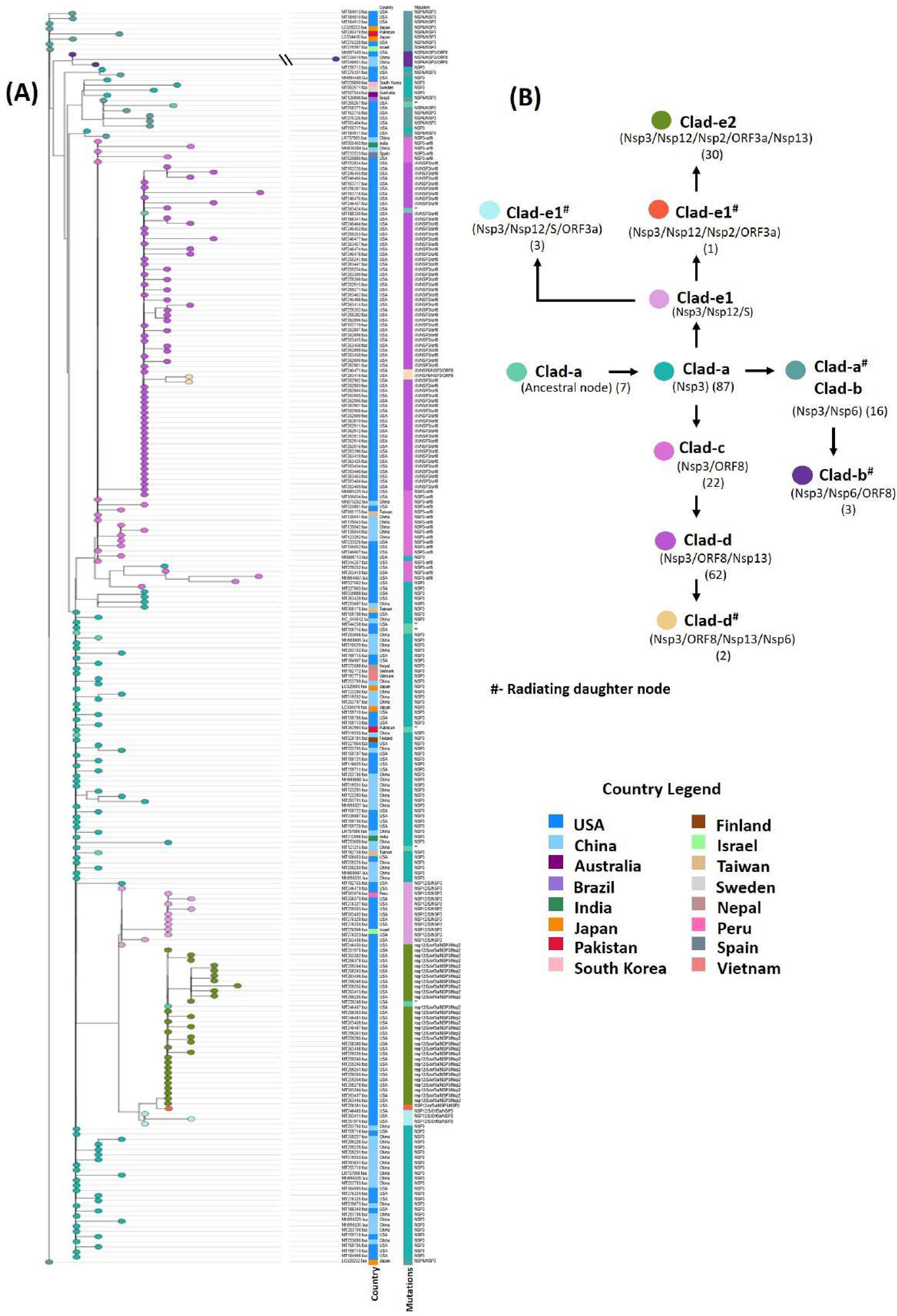
AAV based phylogenetic map of 245 SARS-CoV-2 genomes. Node color represents co-mutational combinations. The formation of each clade is well correlated with the mutational combinations (nG=10).

### The assessment of mutations in SARS-CoV-2 proteins

Amino acid variations were predicted in eight (Nsp2, Nsp3, Nsp6, Nsp12, Nsp13, S, Orf3a, Orf8) SARS-CoV-2 proteins (Table 1). To identify their potential functional role, we carried out the structural analysis of the proteins. Pairwise sequence alignment of wild-type and mutant proteins provided the exact location and changes in amino acids. The GMQE and QMEAN values ranged from 0.45 to 0.72 and −1.43 to −2.81, respectively. The sequence identity ranged from 34% to 99%, which suggested models were constructed with high value of confidence (Figure 4). The I-Mutant DDG tool predicts if a mutation can largely destabilize the protein (ΔΔG<-0.5 Kcal/mol), largely stabilize (ΔΔG>0.5 Kcal/mol) or have a weak effect (−0.5<G=ΔΔG<G=0.5 Kcal/mol). The protein stability analysis showed that all the identified mutations decreased the stability of seven proteins (Nsp2, Nsp6, Nsp12, Nsp13, S, Orf3a, Orf8) except Nsp3 (T1103P) which predicted to increase protein stability (Figure 4 A-H). Further, to explore the role of mutations in SARS-CoV-2 proteins, we carried out HOPE analysis. D614G mutation in S-protein could disturb the rigidity of the protein and due to glycine, hydrophobicity will affect the intra hydrogen bond formation with G594. In ORF8 and Nsp3, the mutation location was not conserved, hence it did not affect or damage the protein function. The mutation (P409L) in Nsp13 was present in the RNA virus helicase C-terminal domain. Since proline is a very rigid amino acid and therefore induce a particular backbone conformation that might be required at this position so this mutation could disturb domain and abolished its function. Mutation L37F (Nsp6) and T85I (Nsp2) were also highly conserved thus could profoundly damage the function of the respective protein. The P324L (Nsp12) mutation was in the RNA binding domain located on the surface of the protein; modification of this residue could disturb interactions with other molecules or other parts of the protein. Conclusively, Nsp3 mutation which appeared in all co-mutation combinations, contributed to increased protein stability among 238 strains could be assigned to their increased pathogenicity. Thus, we attempted to highlight the effects of these mutations in host pathogen interactions.

**Figure 4.**
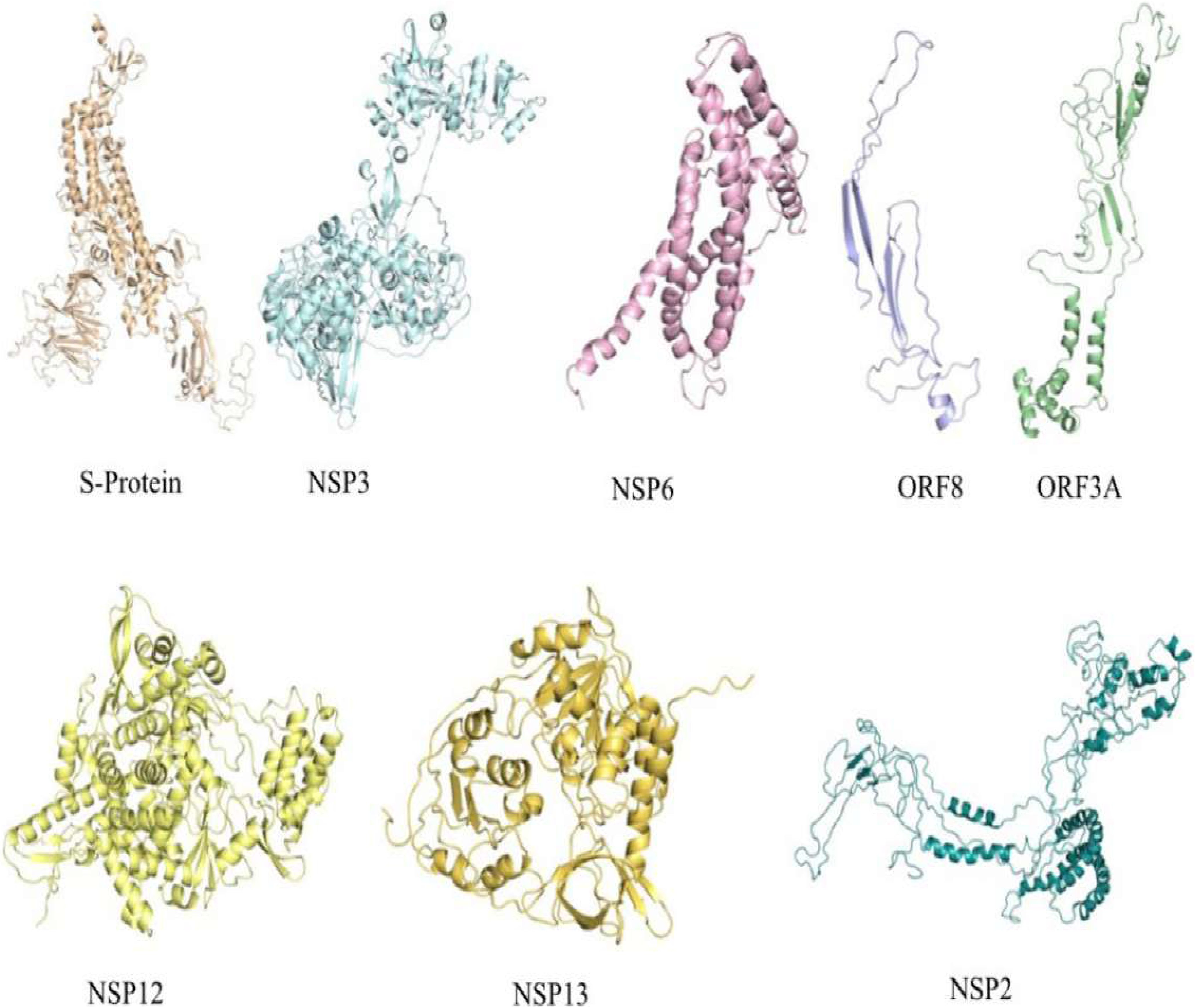
3-D structure prediction of SARS-CoV-2 proteins harboring mutations at different locations to check for its stability in the cell. Structure are predicted using SwissModel and Phyre2 servers.

### Modelling of Host-Pathogen Interaction Network and its Functional Analysis

The HPI Network of SARS-CoV-2 (HPIN-SARS-CoV-2) contained 159 edges, 81 nodes, including 21 viral and 60 host proteins (Figure 5A). The significant existence of few main gene hubs, namely, N, S and M in the network and the attraction of a large number of low-degree nodes toward each hub showed strong evidence of controlling the topological properties of the network by few hubs proteins; N with 37 degrees, S, and M with 17 and 8 degrees, respectively. These viral proteins are the main hubs in the network, which regulate the network. Based on degree distribution, the viral protein N showed highest pathogenicity followed by S and M. N is a highly conserved major structural component of SARS-CoV-2 virion involved in pathogenesis and used as a marker for diagnostic assays (54). Another structural protein S (spike glycoprotein), attach the virion to the cell membrane by interacting with host receptor, initiating the infection (55). The M protein, component of the viral envelope played a central role in virus morphogenesis and assembly via its interactions with other viral proteins (56). Interestingly, we found four host proteins MYO5A, MYO5B, MYO5C and T had a maximum interaction with viral hub proteins. MYO5A, MYO5B, MYO5C interacting with all three (N, S and M) whereas T with two (S and M) viral hub proteins, showed a significant relationship with persistent infections caused by the SARS-CoV-2. Other host proteins showing highest degree namely, ATP6V1G1 and RPS6 were found interacting with all the NSPs and polyprotein of ORF1a respectively.

**Figure 5.**
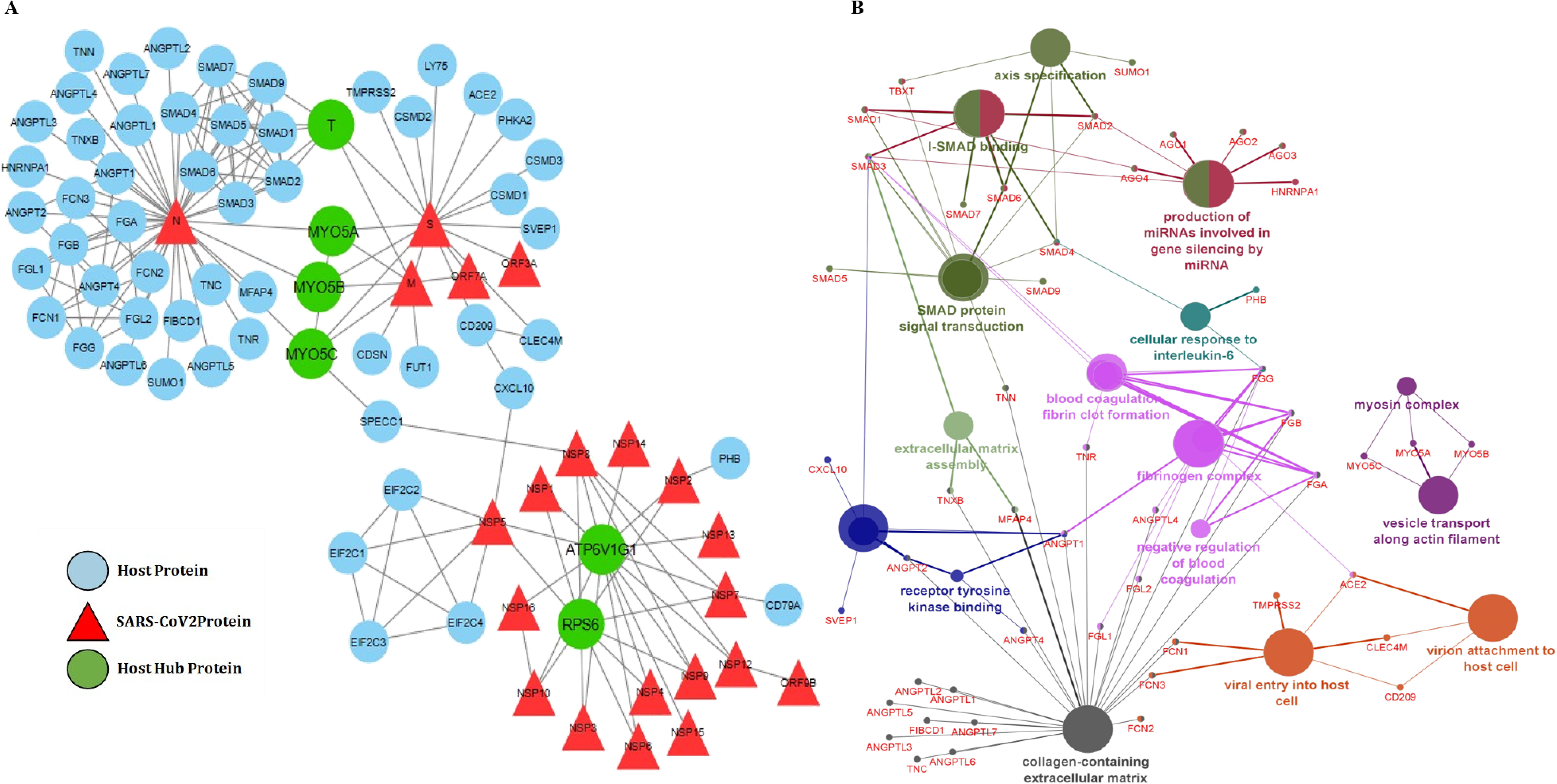
(A) Host-pathogenic interaction of SARS-CoV-2 and human proteins. Nodes represented proteins while lines/edges representing interaction. Triangles (Red) represent viral proteins found to be directly interacting with the human proteins (blue). The hubs (MYO5A, MYO5B, MYO5C, T, RPS6 and ATP6V1G1 (green) were found interacting with maximum viral proteins. (B) Gene ontology (GO) analysis was performed for host proteins using the ClueGo Cytoscape app against database KEGG, the Gene Ontology—biological function database, and Reactome pathways. ClueGo parameters were set as follows: Go Term Fusion selected; P values of ≤0.05; GO tree interval, all levels; kappa score of 0.42.

MYO5A, MYO5B and MYO5C proteins are Class V myosin (myosin-5) molecular motor that functions as an organelle transporter (57, 58). The presence of myosin protein played a crucial role in coronavirus assembly and budding in the infected cells (59). These cytoskeletal proteins are of importance during internalization and subsequent intracellular transport of viral proteins. It was found that inhibition of MYO5A, MYO5B, and MYO5C was efficient in blocking the internalization pathway, thus this target can be used for the development of a new treatment for SARS-CoV-2 (60). Patients suffering from COVID-19 undergo two major condition in the severe stage, thrombotic phenomenon and hypoxia, that are acting as silent killers (61, 62). Hypoxia, condition where oxygen level of the body reduces drastically results in the elevated expression of T protein in the body (63). T protein (Brachyury/TBXT) is transcription factor involved in regulating genes required for mesoderm formation and differentiation thus playing an important role in pathogenesis. ATP6V1G1 (Catalytic subunit of the peripheral V1 complex of vacuolar ATPase) is responsible for acidifying a variety of intracellular compartments in eukaryotic cells. It is reported that Nsp5 may cleave host ATP6V1G1 thereby modifying host vacuoles intracellular pH (64). RPS6 plays an important role in controlling cell growth and proliferation through the selective translation of particular classes of mRNA. Reports have shown downregulation of RPS6 during the infection severity (65). The detailed functional analysis of HPIN-SARS-CoV-2 was mapped on the radiological findings from the COVID-19 severely infected patients and non-survivors. It was reported that the levels of fibrin-degrading proteins, fibrinogen and D-dimer protein were 3-4 folds higher as compared to healthy individual. Therefore, reflecting coagulation activation from infection/sepsis, cytokine storm and impending multiple organs failure (66–69). In our network, we found 47 proteins (SUMO1, T, SMAD1-9, AGO1-4, HNRNPA1, PHB, TNN, TNR, TNXB, CXCL10, SVEP1, ANGPT1-2, ANGPT4, ANGPTL1-7, MYO5A, MYO5B, MYO5C, FGL1-2, FCN1-3, ACE2, TMPRSS2, CLEC4M, CD209, FGA, FGB, FGG) are associated with the above etiology (Figure 5B). We also found the interaction of SMAD family proteins and SUMO1 with N protein, which may result to inhibition of apoptosis of infected lung cells. The interactome study reveals a significant role of identified host proteins in viral budding and related symptoms of COVID-19.

### The mutation in SARS-CoV-2 proteins inhibit viral penetration into host

To validate the effect of amino acid variation (AAV), significant host proteins interactions from HPIN-SARS-CoV-2 were considered for *in silico* docking studies. Docking of S-Protein (wild type and mutant) with ACE2, TMPRSS2 and one of myosin proteins (MYO5C) were analyzed. Recent studies have shown that SARS-CoV-2 uses the ACE2 for entry and the serine protease TMPRSS2 for S protein priming (70). The polyproteins (Nsp12, Nsp13, Nsp2, Nsp3 and Nsp6) of ORF1A and ORF1AB were docked with RPS6 and ATP6V1G1 host proteins. The docking results showed that mutant S-protein could not bind efficiently with ACE2 and MYO5C, whereas mutation slightly promotes the binding with TMPRSS2 (Table 3, Figure 6 and Figure 5B). TMPRSS2 has been detected in both nasal and bronchial epithelium by immunohistochemistry (71), reported to occur largely in alveolar epithelial type II cells which are central to SARS-CoV-2 pathogenesis (72). The wild-type S-protein form 16 hydrogen bonds and 1058 non-bonded contacts with ACE2; whereas the mutant protein forms 12 hydrogen bond and 738 non-bonded contacts (Figure 6). This result suggests that D614G mutation in S-protein could affect viral entry into the host. Similarly, mutations present in the Nsp12, Nsp13, Nsp2, Nsp3 and Nsp6 of SARS-CoV-2 could inhibit the interaction with RPS6, but these mutations promote the binding with ATP6V1G1 expect Nsp6 (L37F). The RPS6 contributes to control cell growth and proliferation (73), so a loss of interaction with RPS6 could probably inhibit the production of viruses. Overall structural and interactome analyses suggests that identified mutations (Nsp2 (T85I), Nsp3 (S1103P), Nsp6 (L37F), Nsp12 (P324L), Nsp13 (P409L, Y446C), S (D614G)) in SARS-CoV-2 might play an important role in modifying the efficacy of viral entry and its pathogenesis. However, these observations required critical revaluation as well as experimental work to confirm the *in-silico* results.

**Table 3.**
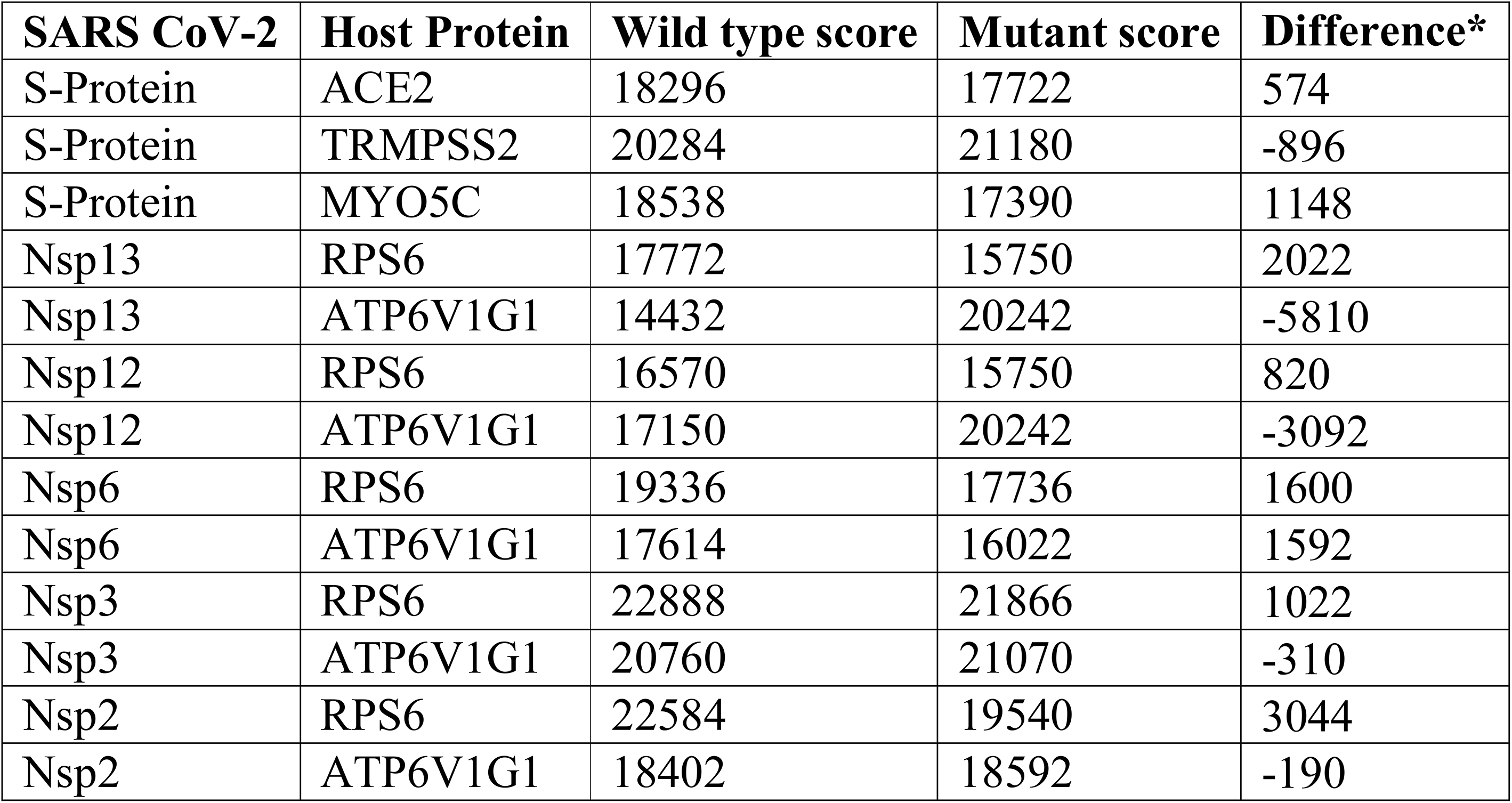
*In silico* docking analysis of SARS-CoV-2 proteins with Human proteins

**Figure 6.**
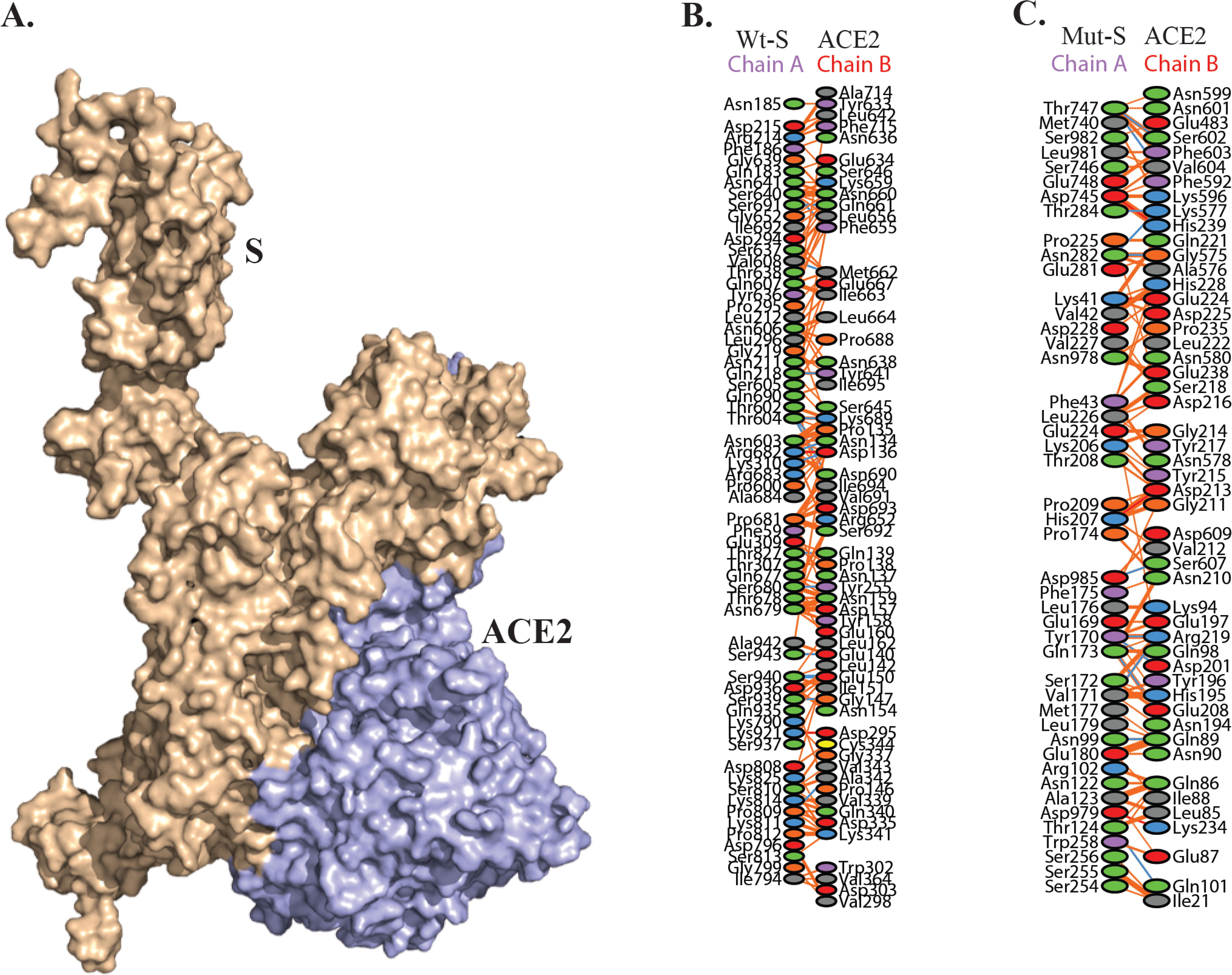
*In-silico* receptor-ligand docking analysis for mutated S protein (D614G) from SARS-CoV-2 and ACE2 protein present in human. B & C represents amino-acid interactions between wild type and mutated Spike protein with ACE2 receptor.

### Regulation of SARS-CoV-2 pathogenicity by CpG island

The genotyping analysis that we performed showed high frequency rate (45) of SNPs at 5’UTR region (Table 1) and recent study also suggested that suppression of GC content could play a vital role in specific antiviral activities (54). As seen in SNP analysis, the common transitions of C>T and G>A that alters the GC content of the SARS-CoV-2 (Table 1), directed the prediction of CpG dinucleotides which are involved in silencing of transcription and down regulation of viral replication (74). In RNA viruses, CpG dinucleotides are targeted by Zinc Antiviral protein (ZAP), an intracellular broad-spectrum antiviral restriction factor which plays a vital role in generating innate immune response against wide range of RNA viruses in vertebrates (75, 76). ZAP mediated Antiviral Restriction has been already demonstrated against different RNA viruses including Flaviviruses, Filoviruses, Influenza, Alphaviruses and Retroviruses (77–83). ZAP directly binds to viral RNA through CCCH (Cys-Cys-Cys-His) type zinc finger motifs present at the N-terminal and recruits RNA processing exosome for viral RNA degradation (75, 84). In association with TRIM25, ZAP binds specifically to viral RNA regions with elevated CpG dinucleotide frequencies, leading to inhibition of replication and translation of viral RNA (14, 85–87).

Thus, CpG dinucleotide motif profiling and their importance of existence in SARS-CoV-2 genomes was proceeded. We found that CpG islands were consistently present in two regions of the genome at the positions 285-385 nucleotides (101 bp) and 28,324-28,425 nucleotides (102 bp). The results were consistent in all 245 genomes analyzed in the present study with 100% conservancy in 237 genome sequences (Figure 7).

**Figure 7.**
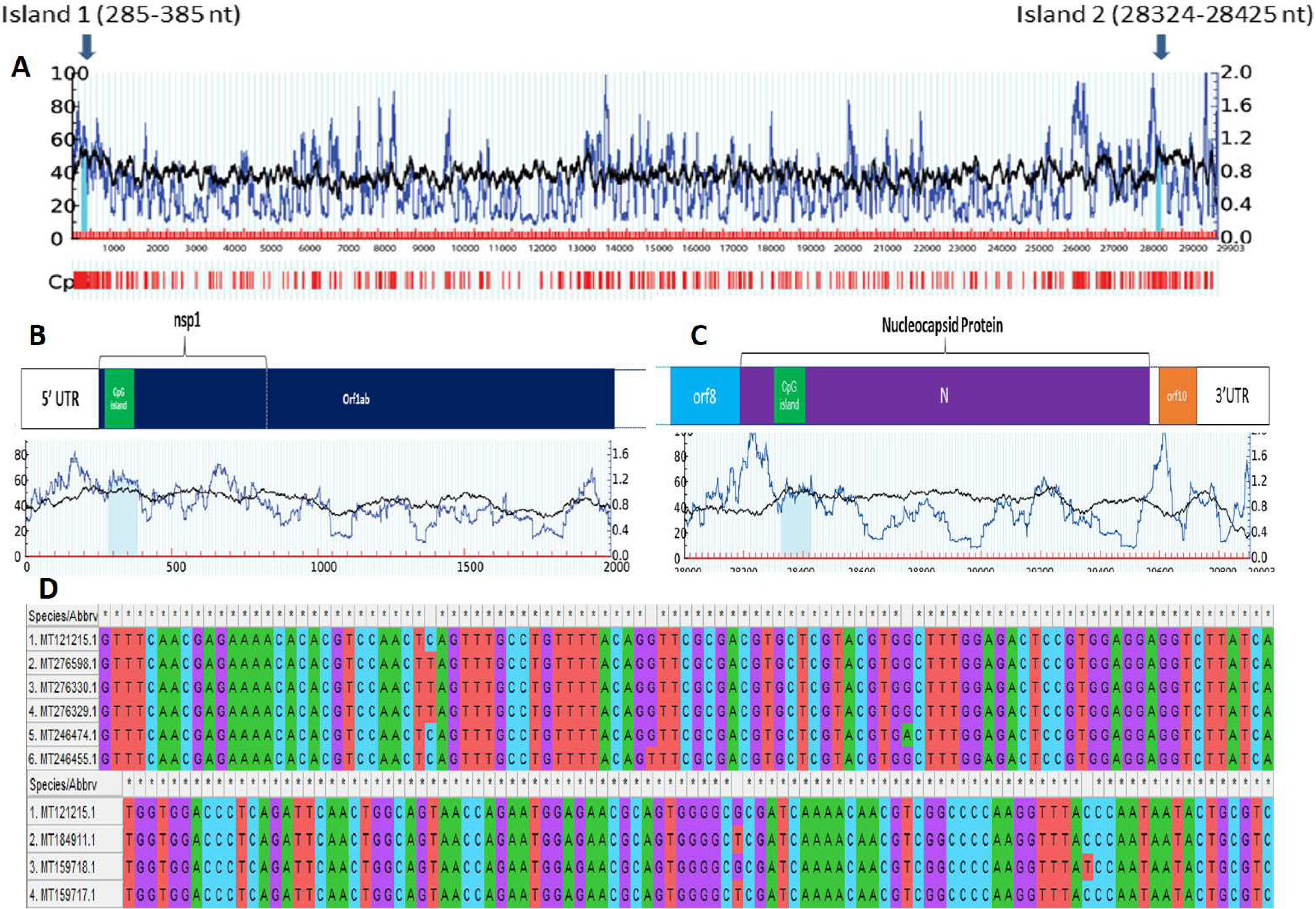
Detection of two CpG islands in Wuhan_Hu-1 complete genome sequence (Accession number: MT121215.1), marked by blue arrows. One of the CpG island was found to be located towards the 5’ end of the genome, in ORF1ab. Another CpG island was found towards the 3’ end of the genome, located in ORF9 coding for N protein.

In the remaining 8 genomes, 5 genomes (MT246474.1 (G to A substitution at 354^th^ position with respect to reference genome); MT276329.1, MT276330.1 and MT276598.1 (C to T substitution at 313^th^ position) and MT246455.1 (G to T substitution at 332^nd^ position)) showed point mutation in 5’ CpG island; whereas three genomes (MT159718.1 (C to T substitution at 28409^th^ position); MT159717.1 and MT184911.1 (G to T substitution at 28378^th^ position)) showed point mutation in 3’CpG end. Interestingly, all these sequences belong to USA. On further locating CpG island positions with respect to proteins, it was found that these two CpG islands were located at two prime locations within the genome, one in Nsp1, and another within N protein. Previously, it was reported that both the proteins interacted with 5’ UTR region playing crucial roles in viral replication and gene expressions (4, 88, 89). Most pivotal role of N protein revolves around encapsulation of viral gRNA which leads to formation of ribonucleoprotein complex (RNP), which is a vital step in assembly of viral particles (90).

Nsp1 protein in coronaviruses plays a regulatory role in transcription and viral replication (90). It is known to interact with 5’ UTR of host cell mRNA to induce its endonucleolytic cleavage (91, 92), thus inhibiting host gene expression (93). It also plays an important role in blocking IFN-dependent antiviral signaling pathways leading to dysregulation of host immune system (94–96). CpG sites can be targeted by Zinc Finger Antiviral Proteins which can mediate antiviral restriction through CpG motif detection (77, 82, 83). Apart from this, CpG oligodeoxynucleotides (ODNs) are known to act as adjuvants and are already established as a potent stimulator for host immune system (97–100). Moreover, recent studies conducted on influenza A and Zika virus genome has shown that by increasing the CpG dinucleotides in viral genome, impairment of viral infection is observed (101, 102). Our result showed that the presence of conserved CpG islands in Nsp1 and N protein across all genomes of SARS-CoV-2 indicated their role in pathogenesis and can be targeted by Zinc Finger Antiviral Proteins or exploited to design CpG-recoded vaccines.

## Conclusions

The genomic and proteomic survey of SARS-CoV-2 strains reported from subset of population of different countries reflected global transmission during the outbreak of COVID-19. The viral phylogenetic network with five clades (a-e) provided a landscape of the current stage of epidemic where major divergence was observed in USA strains. From this we propose genotypes linked to geographic clades in which signature SNPs can be used to track and monitor the epidemic. Demarcation of co-mutation in the SARS-CoV-2 strains by assessing co-mutations also highlighted the evolutionary relationships among the viral proteins. Our results suggested that co-mutations are indicative of AAV based induced pathogenicity leading to multiple mutations embedded in few genomes. It was also seen that just increasing the genomic sample size by 50 times did not led to prediction of significant mutations or co-mutations that were leading to strain variation in SARS-CoV-2 virus. Thus, sample size of SARS Cov-2 genome does not have a direct relation with variation to be predicted in amino acid. However, co-mutations are still in evolutionary process and more combinations can be predicted with a large dataset. High-frequency AAV mutations were present in the critical proteins, including the Nsp2, Nsp3, Nsp6, Nsp12, Nsp13, S, Orf3a, Orf8 which could be considered for designing a vaccine. Comparative analysis of proteins from wild and mutated strains showed positive selection of mutation in Nsp3 but not in rest of the mutants. The HPI model can be used as the fundamental basis for structure-guided pathogenesis process inside host cell. The interactome study showed MYO-5 proteins as a key host partner and highlighted the key role of N, S and M viral proteins for conferring SARS-CoV-2 pathogenicity. The mutation in the S protein could affects the viral entry by loose binding with ACE2. The presence of CpG dinucleotides in N and Nsp1 protein could play a critical role in pathogenesis regulation. Based on our multi-omics approach: genomics, proteomics, interactomics, systems and structural biology provided an opportunity for better understanding of COVID-19 strains and its mutational variants.

## Supporting information

Data Set S2

Data Set S3

Data Set S1

## Acknowledgements

VG acknowledges Phixgen Pvt. Ltd. for research fellowship. MV, SS acknowledge Dr. P. Hemalatha Reddy, Principal, Sri Venkateswara College, University of Delhi for her constant support and encouragement. RL and US also acknowledge The National Academy of Sciences, India, for support under the NASI-Senior Scientist Platinum Jubilee Fellowship Scheme. NS acknowledges Council of Scientific and Industrial Research (CSIR), New Delhi for doctoral fellowships. HV would like to thank Ramjas College, University of Delhi, Delhi for providing support. RK acknowledges Magadh University, Bodh Gaya for providing support. PH would like to thank Maitreyi College, University of Delhi, Delhi for providing support. YS acknowdge J.C. Bose (SERB) fellowship.

---

**Supplementary Figure 1.** AAV based phylogenetic map of 12299 SARS-CoV-2 genomes. The formation of each clade is well correlated with the mutational combinations produced by 9 differenet mutation. The green clade is represented by the 245 genomes and addition of ORF7, N, nsp2, Orf3a mutation to Clad e1 resulted in delineation of other combinations to form clad-f-j.

## Supplementary Data

**Data Set S1**: Summary of the SARS CoV-2 genomes with isolated geographical location selected for comparative genomics analysis. Sheet1: 447 Genomes, Sheet2: 245 Genomes, Sheet: 18775 genomes & Sheet4: 12299 Genomes.

**Data Set S2:** Summary of the mutations identified from the 245 SARS-CoV-2 genomes and selected co-mutations.

**Data Set S3:** Summary of the mutations identified from the 12299 SARS-CoV-2 genomes.

