## Supplementary figures and images for "Comparative Genomics and Integrated Network Approach Unveiled Undirected Phylogeny Patterns, Co-mutational Hotspots, Functional Crosstalk and Regulatory Interactions in SARS-CoV-2"

### Supplementary Figure 1

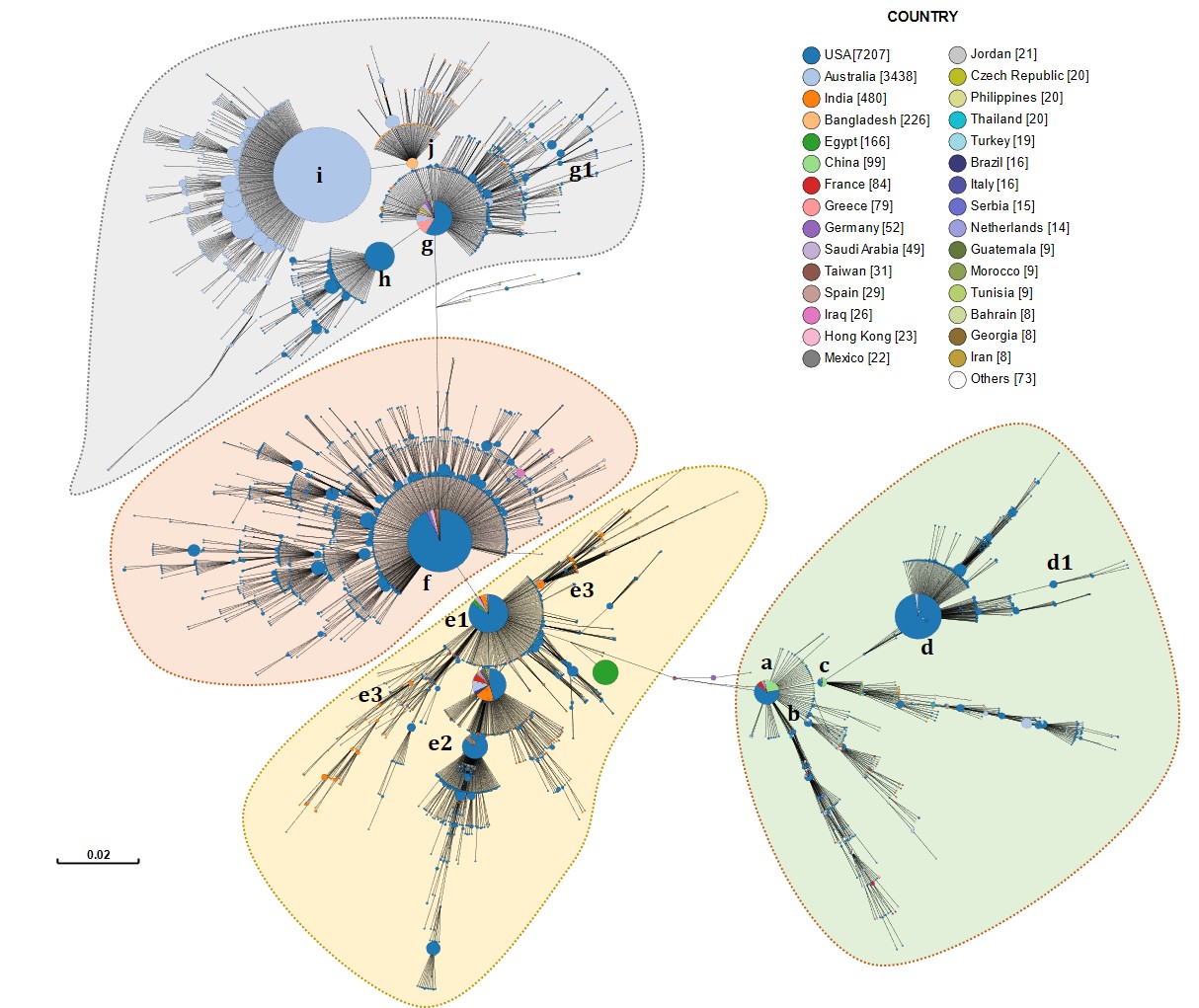
